# Immune regeneration in irradiated mice is not impaired by the absence of DPP9 enzymatic activity

**DOI:** 10.1101/431775

**Authors:** Margaret G. Gall, Hui Emma Zhang, Adam Cook, Christopher J. Jolly, Geoffrey W. McCaughan, Ben Roediger, Mark D. Gorrell

**Author notes:** Equal first author (Margaret G. Gall, Hui Emma Zhang). **Corresponding author:** Mark D. Gorrell, Liver Enzymes in Metabolism and Inflammation Program, Centenary Institute, Locked Bag No. 6, Newtown, NSW 2042, Australia.

## Abstract

The ubiquitous intracellular protease dipeptidyl peptidase 9 (DPP9) has roles in antigen presentation and B cell signaling. To investigate the importance of DPP9 in immune regeneration, primary and secondary chimeric mice were created in irradiated recipients using fetal liver cells and adult bone marrow cells, respectively, using wild-type (WT) and DPP9 gene-knockin (DPP9^S729A^) enzyme-inactive mice. Immune cell reconstitution was assessed at 6 and 16 weeks post-transplant. Primary chimeric mice successfully regenerated neutrophils, natural killer, T and B cells, irrespective of donor cell genotype. There were no significant differences in total myeloid cell or neutrophil numbers between DPP9-WT and DPP9^S729A^-reconstituted mice. There were fewer B cells in the DPP9^S729A^-origin compared to DPP9 WT-origin mice in primary chimeric mice. In secondary chimeric mice, cells of DPP9^S729A^-origin cells displayed enhanced engraftment compared to WT. However, we observed no differences in myeloid or lymphoid lineage reconstitution between WT and DPP9^S729A^ donors, indicating that hematopoietic stem cell (HSC) engraftment and self-renewal is not diminished by the absence of DPP9 enzymatic activity. This is the first report on transplantation of bone marrow cells that lack DPP9 enzymatic activity.

## Introduction

The ubiquitous intracellular post-proline serine protease dipeptidyl peptidase 9 (DPP9) belongs to the DPP4 gene family, which includes four atypical serine proteases: DPP4, fibroblast activation protein (FAP), DPP8 and DPP9 ^1,2^. DPP9 plays roles in both innate and adaptive immunity. DPP9 is extensively expressed throughout immunological tissues *in vivo* ^3^ and within individual leukocyte subpopulations ^1,4–9^. DPP9 mRNA and protein is up-regulated in stimulated mouse splenocytes and in Jurkat T- and Raji B-cell lines ^6^. Endogenous DPP9 limits the presentation of an antigenic peptide, RU134–42, through cleaving this peptide ^10^. DPP9 causes Syk degradation and thus influences Syk signalling in B cells ^8^. Activation and proliferation of innate and adaptive immune cells is diminished in the absence of DPP9 enzymatic activity ^4,9,11,12^. Within monocytes and macrophages, basal DPP8 and DPP9 activity suppresses inflammasome activation through inhibition of pro-caspase-1 activation via NLRP-1 ^13,14^. Thus, a variety of evidence supports multiple roles for DPP9 in the regulation of immune function.

We generated the first gene DPP9 knock-in (DPP9^S729A^) mouse that has a single serine-to-alanine point mutation at the enzyme active site (S729A) ^15^. Unlike mice deficient in any other protease of this gene family, homozygote DPP9 deficiency is neonate lethal ^15–17^. DPP9 is closely related to the extracellular proteases DPP4 (CD26) and fibroblast activation protein (FAP) ^18^. DPP4 is expressed by immune cells of both the myeloid and lymphoid lineages ^19 20^. Genetic or pharmacologic ablation of DPP4 improves bone marrow engraftment ^21^. We found that FAP expression does not influence the proportions of CD4+ and CD8+ T cells, B cells, dendritic cells and neutrophils in the thymus, lymph node or spleen in healthy adult mice ^22^. Whether the absence of DPP9 enzymatic activity affects short-term and long-term repopulation of immune cells of the lymphoid or myeloid lineages is underexplored.

Hematopoiesis is critically dependent upon hematopoietic stem cells (HSC). HSC migrate into the fetal liver between embryonic day (ED) 11 and 12 whereupon their numbers expand substantially ^23,24^. Between ED 13.5 and 14.5, the fetal liver contains large numbers of hematopoietic foci with erythropoiesis constituting a major part of their activity but also with capacity for myelopoiesis and lymphopoiesis ^25^. A successful short-term primary engraftment (30 to 60 days) can provide confirmation that the progenitor cell pool is intact and that all myeloid and lymphoid cell types are present and, in the long term (4 months) whether the reconstituted HSC are functional ^26–28^. However, even successful long-term engraftment in a primary transplant recipient does not rule out defects in self-renewal or proliferation capability. Hence, a further serial transplant is often undertaken in chimera studies to demonstrate intact HSC engraftment and renewal ^27^.

Post-transplant, identifying the progeny of the transplanted HSC is important to ascertain the effectiveness of the original graft and the properties of the regenerating immune system. The most commonly used method to achieve this is through the CD45 allelic model, where genetic differences in CD45 (CD45.1 and CD45.2) between donor and recipient mouse strains enable donor-derived cells to be traced by flow cytometry ^26,29^. Neutrophils and macrophages are the first cell types to recover after combined myelo-ablative irradiation and fetal liver or adult bone marrow cell transplant. These cells appear in the first few days after transplant, followed closely by B cells. Platelets and red blood cell lineages are present in the peripheral circulation at one to two weeks post-irradiation ^27^. A small proportion of host T cells resist the effects of irradiation and expand in the post-irradiated environment, and can be detected within three weeks of transplant, while donor T cells usually become detectable 4 to 5 weeks after transplantation ^29^.

Very recently, an independent study found that ED 17.5 fetal liver-derived hematopoietic stem cells from a similar DPP9^S729A^ mouse ^16,17^ are able to fully reconstitute immune cell subsets 6 weeks after transplant in competitive mixed chimeras ^30^. Here, we have explored the role of DPP9 enzyme activity in immune cell development through the creation of two sequential chimeras using ED 13.5 to 14.5 fetal liver cells and using adult bone marrow cells of WT and DPP9^S729A^ origin. Both short-term (6 weeks) and long-term (4 months) hematopoietic regeneration were analyzed.

## Materials and Methods

### Ethics statement

All animal handling and experimental procedures were approved by Sydney Local Health District Animal Welfare Committee under ethics protocol 2013/017 (K75/5-2012/3/5754), and conducted in accordance with all applicable laws and guidelines. A body scoring system was used recording details for each mouse of posture, activity and gait, breathing, hydration, the presence of abnormal excreta, body condition and body weight as compared to the baseline weight. A weight loss of greater than 15% compared to original weight triggered euthanasia of the mouse.

### Mice

Mice were maintained in the Centenary Institute animal facility under specific pathogen free conditions and exposed to a 12 h light-dark cycle. C57BL/6J and PTPRCA mice were purchased from either Animal Resource Centre (Perth, WA, Australia) or Australian BioResources (Moss Vale, NSW, Australia). Mice were housed 4–6 mice per cage with ad libitum access to rodent chow (Specialty Feeds, Perth, West Australia) and water and entered experiments more than one week after any transport. All the mice used for chimera experiments were at least 8 weeks old before irradiation.

The DPP9^S729A^ mouse strain has been described ^15^. All embryos required for fetal liver cells were obtained from pregnant females where pregnancy resulted from timed mating of DPP9 heterozygous intercrosses. For mating, single females were placed in the male home cage for 12 h then checked for the presence of a vaginal plug as an indicator of mating prior to separation from the male. The time of separation of female and male was designated ED 0.5. Females were weighed at ED 0.5 and weighed daily from ED 4.5 until ED 13. Pregnancy was suggested by weight gain greater than non-pregnant between ED 7.5 and ED 13.5 ^31^.

### Generation of fetal liver chimeras

Primary chimeras were generated using donor-derived fetal liver cells ^26^. Liver cells from embryonic mouse livers at ED13.5 – ED14.5 from DPP9^S729A^ and DPP9-wildtype (DPP9-WT) pups were excised and mashed through a 70 μm sieve into liver cell buffer (1% Glutamax, 2% Penistrep, 10% FCS in Dulbecco’s Modified Eagle’s Medium). The cell suspension was centrifuged at 2000 rpm for 5 min and the pellet resuspended in cell freezing solution (10% DMSO in FCS). It was necessary to store liver cells by freezing until genotyping results confirmed. When thawing, cell suspension was centrifuged at 2000 rpm for 7 min and the pellet resuspended in 10 mL liver cell buffer. After standing at room temperature for 30 min to release the maximum amount of DMSO, the cell suspension was centrifuged at 4°C at 2000 rpm for 7 min and resuspended in 400 μL of Hank’s balance salt solution (HBSS) for injection. Eighteen 10 week-old male PTPRC^A^ mice (carrying the CD45.1 allele) received two doses of a 600 rad exposure (at a dose rate of 110 rad per minute) of myelo-ablative radiation using a Gammacell^®^ 40 Exactor Low Dose-Rate Research Irradiator (MDS Nordion Inc. Ontario, Canada) administered 4 h apart as previously described ^32^. For fetal liver cell transplant, 2 × 10^6^ cells at a volume of 200 μL per recipient mouse was injected via the lateral tail vein into recipient mice 3–4 h after irradiation. Post-transplantation, mice were provided with antibiotic in water for 14 days and monitored daily.

### Generation of bone marrow chimeras

Bone marrow (BM) was prepared and transplanted as previously ^32^. Secondary chimeras were produced by transferring regenerated BM donor cells from adult primary chimeric mice into new irradiated recipient mice. Nineteen 10 week-old male PTPRCA mice received one dose of a 750 rad exposure of myelo-ablative radiation as described above. The donors were two healthy DPP9^S729A^-origin primary chimeric mice and two DPP9-WT-origin primary chimeric mice (all carrying the CD45.2 allele). Each pair of selected donor mice had shared a cage but the chimerism in each mouse was produced with a different batch of fetal liver cells. BM cells from these four mice, along with two control male PTPRCA mice, were harvested. BM were released from the femur, tibia and fibula, rinsed and filtered through a 70 μm and then a 40 μm filter. Cell suspension was centrifuged at 300 × g for 5 min at 4°C and the pellet re-suspended in HBSS. For bone marrow transplant, 6.7 × 10^6^ primary chimera bone marrow cells and 10 × 10^6^ competitor bone marrow cells (carrying the CD45.1 allele) from PTPRCA mice (ratio of donor: competitor = 40%: 60%) at a volume of 200 μL per recipient mouse were injected via the lateral tail vein into recipient mice 3–4 h after irradiation. Post-transplantation, mice were provided with antibiotic water for 14 days and monitored daily.

### Flow cytometry

For analysis of fetal liver cells, cell aliquots were thawed and red blood cells were lysed (8.26 g ammonium chloride, 1 g potassium bicarbonate and 0.037 g EDTA dissolved in 1 L of dH_2_O). After centrifugation and washing, cell pellet was resuspended in 200 μL of FACS wash. Then 100 μL of stain cocktail containing TER-119-APC and CD11b-FITC antibodies (Table 1) was added and incubated on ice for 1 h. For analysis of peripheral blood, 5–6 drops of blood from tail vein were collected into an Eppendorf containing 50 μL 0.1M EDTA. After red blood cell lysis, cells were pelleted and re-suspended in 200 μL FACS wash. Then 100 μL of stain cocktail containing a number of antibodies to identify immune cell populations represented in a reconstituted immune system (Table 1) was added and incubated on ice for 1 h. Just prior to flow analysis, cells were filtered and DAPI was added (100 ng/mL). Flow cytometry data were acquired using a BD LSR Fortessa cytometer (BD Pharmingen) and subsequently analyzed with FlowJo 10 software (Treestar Inc., Ashland, OR, USA). Gating strategies are shown in Supplementary Figures 1–4. Data were then analyzed by non-parametric Mann-Whitney U test using GraphPad Prism (GraphPad v. 9.9, San Diego, CA, USA).

**Table 1:** Antibodies and their target cell types.

| **Antibody** | **Synonym/**<br>**clone** | **Target cell** | **Dilution**<br>**used** | **Source: catalog #** |
| --- | --- | --- | --- | --- |
| TER-119-APC | TER-119 | Leukocytes of<br>erythroid lineage | 1:250 | BD Biosciences, #557909 |
| CD11b-FITC | M1/70 | Leukocytes of<br>myeloid and NK<br>lineages | 1:250 | BD Biosciences, #557396 |
| BUV395<br>CD19 | 1D3 | B cells | 1:200 | BD Biosciences #563557 |
| BV421 CD3e | 145-2C11 | T cells, NK cells | 1:250 | BD Biosciences #562600 |
| V500<br>CD45.2 | 104 | Leukocytes | 1:250 | BD Biosciences #562129 |
| FITC CD11b | M1/70 | Myeloid cells | 1:250 | BD Biosciences #557396 |
| PerCP-Cy5.5<br>CD45.1 | A20 | Leukocytes | 1:250 | eBioscience<br>#45-0453 |
| PE NK1.1 | PK136 | NK cells | 1:250 | BD Biosciences #557391 |
| AF647 Ly6G | 1A8 | Neutrophils | 1:250 | Biolegend<br>#127610 |
| APC.Cy7<br>CD8a | 53-6.7 | T cells | 1:250 | BD Biosciences #557654 |

## Results

### Hematopoietic stem cells in the fetal liver are unaltered by DPP9 genotype

Fetal liver cells were used as donor cells for primary bone marrow reconstitution chimeras because DPP9^S729A^ mice die within 24 h of birth ^15–17^, precluding the availability of an adult source of DPP9-defective HSC. Prior to chimera experiments, fetal liver cells deficient in DPP9 enzymatic activity were analyzed for viability and the phenotypic content of stem and progenitor cells, which are necessary for immune regeneration. At this developmental stage the majority of haematopoietic cells are committed to the erythroid lineage as indicated by TER-119 reactivity ^33,34^ while the remaining cells predominantly express the myeloid-associated marker CD11b ^35^. Fetal liver cells were harvested at ED13.5 – 14.5 and stained with antibodies against TER-119 and CD11b to identify erythroid and myeloid cells, respectively. The expected predominance of TER-119 positive cells (80–90%) was observed. TER-119 and CD11b immunopositivity was comparable between DPP9-WT and DPP9^S729A^ fetal liver cells (Figure 1).

**Figure 1:**
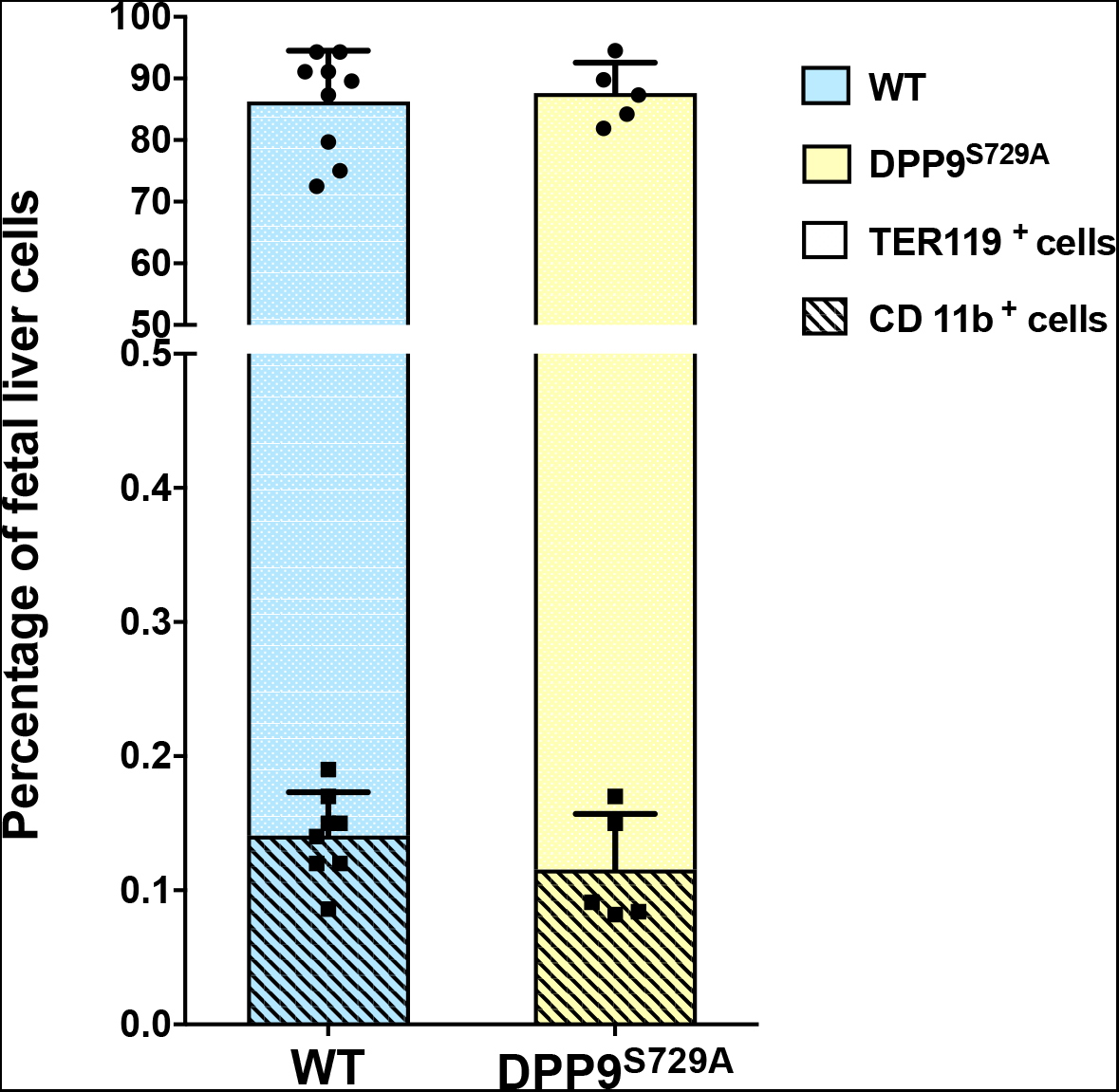
Percentages of TER119^+^ and CD11b^+^ fetal liver cell populations prepared for inoculation of irradiated mice. Fetal liver cells from dPp9^S729A^ and DPP9-WT (WT) embryos were stained with TER119-APC and CD11b-FITC antibodies and analyzed by flow cytometry. No statistically significant difference was observed between the percentages of DpP9^S729A^ and WT cells in the inoculum for either erythroid lineage or myeloid lineage. Stacked bars and error bars represent mean and SD. Black circle and square represent individual data of TER119^+^ and CD11b+ cells, respectively. n = 9 (DPP9-WT embryos) or n = 5 (DPP9^S729A^ embryos).

### DPP9^S729A^ fetal liver cells reconstitute the immune system of irradiated mice

Primary chimeras were generated using fetal liver cells from DPP9-WT and DPP9^S729A^ mice. Peripheral blood was taken at 6 and 16 weeks post-transplantation for flow cytometry analyses of peripheral blood leukocytes, representative of short-term and long-term immune compartment regeneration, respectively.

#### Donor cell survival and hematopoietic replenishment in primary chimeric mice

Mice transplanted with DPP9-WT or DPP9^S729A^ fetal liver cells showed no significant differences in either mean weight or mean percentage weight change throughout the experimental period (Supplementary Figure 5). The replenishment by donor-derived cells in peripheral blood was examined by distinguishing the CD45 alleles in the recipient mice. The proportion of residual recipient cells (CD45.1+) present at 6 weeks post-transplantation was less than 7% of total peripheral blood leukocytes (Figure 2), indicating that the donor cells (CD45.2+) survived the engraftment procedure and that the fetal liver cell transplant conferred a sufficiently effective donor graft to ensure hematopoietic cell reconstitution. At 16 weeks post-transplantation, the residual recipient cells had fallen to ~2% (Figure 2), suggesting effective long-term engraftment by the donor cells. No statistical difference was seen between DPP9-WT and DPP9^S729A^ donor cells at either 6 or 16 weeks post-transplantation. Thus, fetal liver cells carrying the DPP9^S729A^ mutant allele were not impaired in their ability to engraft in irradiated congenic hosts.

**Figure 2:**
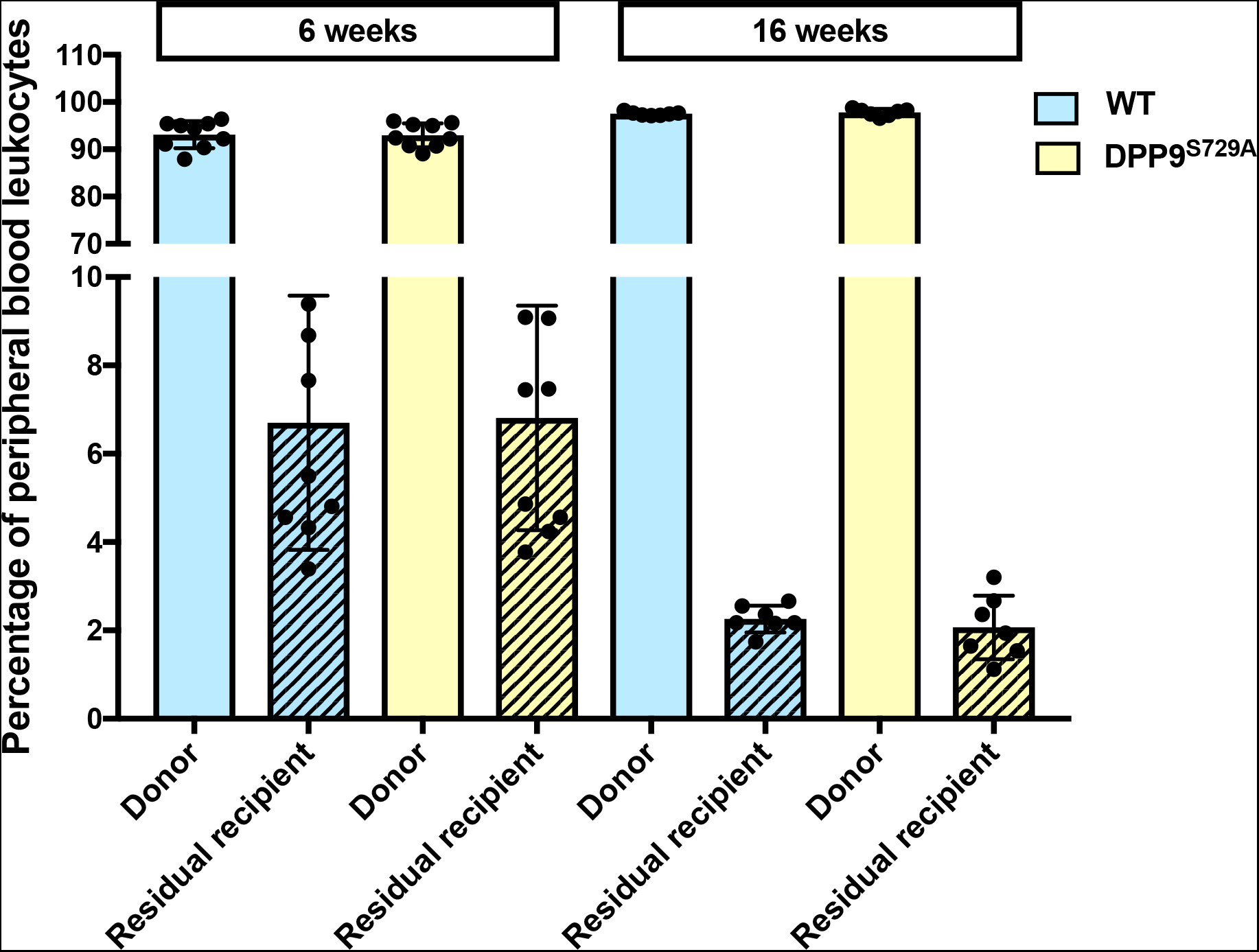
Proportions of donor and residual recipient cells of primary chimeric mice. Irradiated mice inoculated with DPP9^S729A^ or DPP9-WT (WT) fetal liver cells were assessed for the percentage of donor cell-origin and residual recipient cell-origin leukocytes at 6 and 16 weeks post-transplantation. Both DPP9^S729A^ and WT cell-origin groups displayed a small percentage of residual recipient cells at 6 weeks post-transplantation, and fewer at 16 weeks post-transplantation. Mean, SD and individual data. n = 9 (6 weeks) or n = 7 (16 weeks).

#### Identification of myeloid cell types in primary chimeric mice

To examine the development and repopulation of hematopoietic lineages, the major myeloid and lymphoid cell types were analyzed. Comparison was made between DPP9-WT-origin and DPP9^S729A^-origin chimeras as well as normal peripheral blood leukocytes data for C57BL/6 mice ^36^. Here we assessed two major lineages: Ly6G+ neutrophils and CD3− CD19− NK1.1− Ly6G− CD11b+ myeloid cells (which are predominantly monocytes; data not shown). There were no significant differences observed between the WT and DPP9^S729A^ cell-origin recipient mice for either time-point for either total myeloid cells or neutrophils (Figure 3A), suggesting that carrying the mutant DPP9 allele does not significantly alter hematopoietic regeneration in primary chimeric mice.

**Figure 3:**
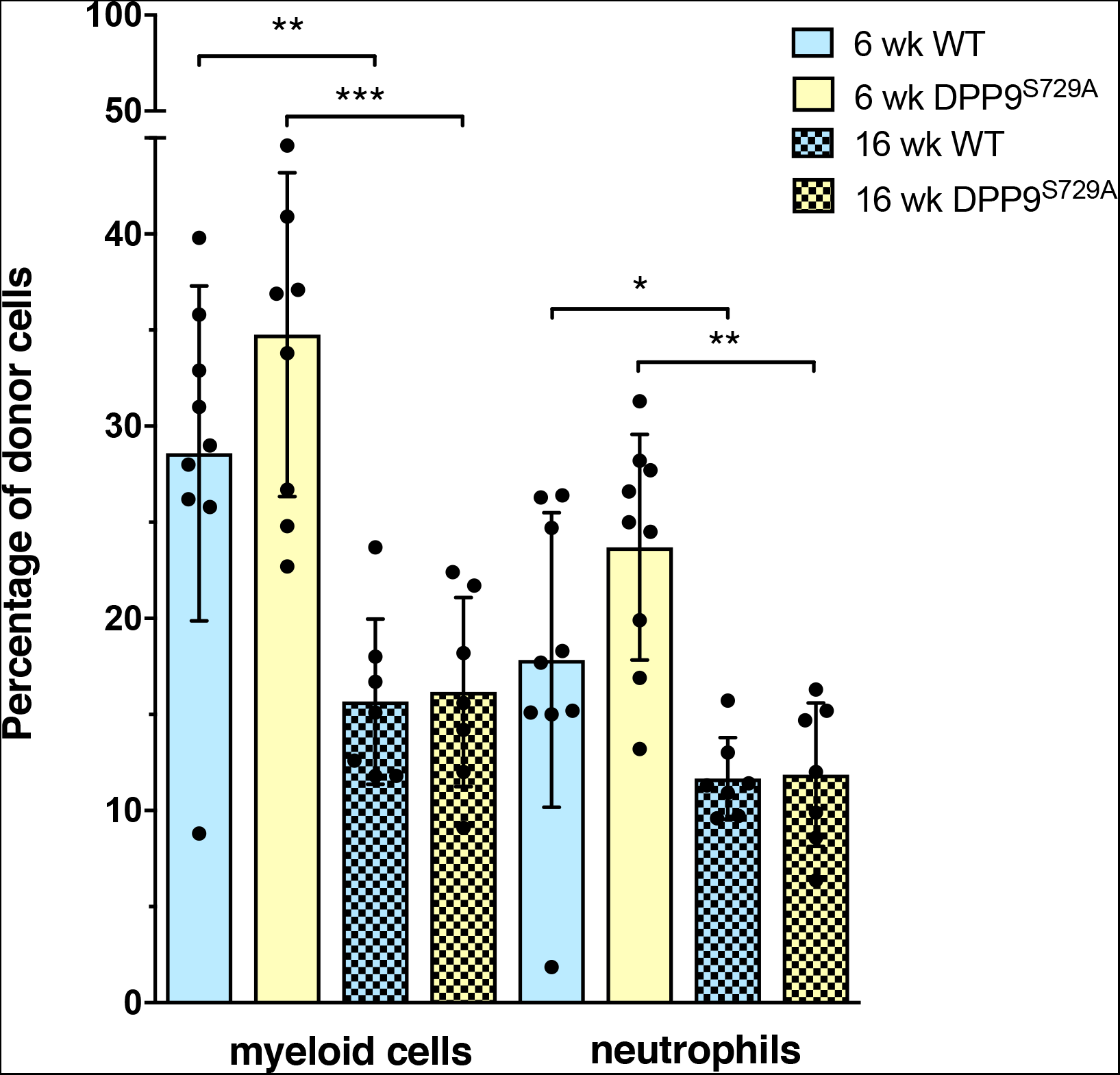
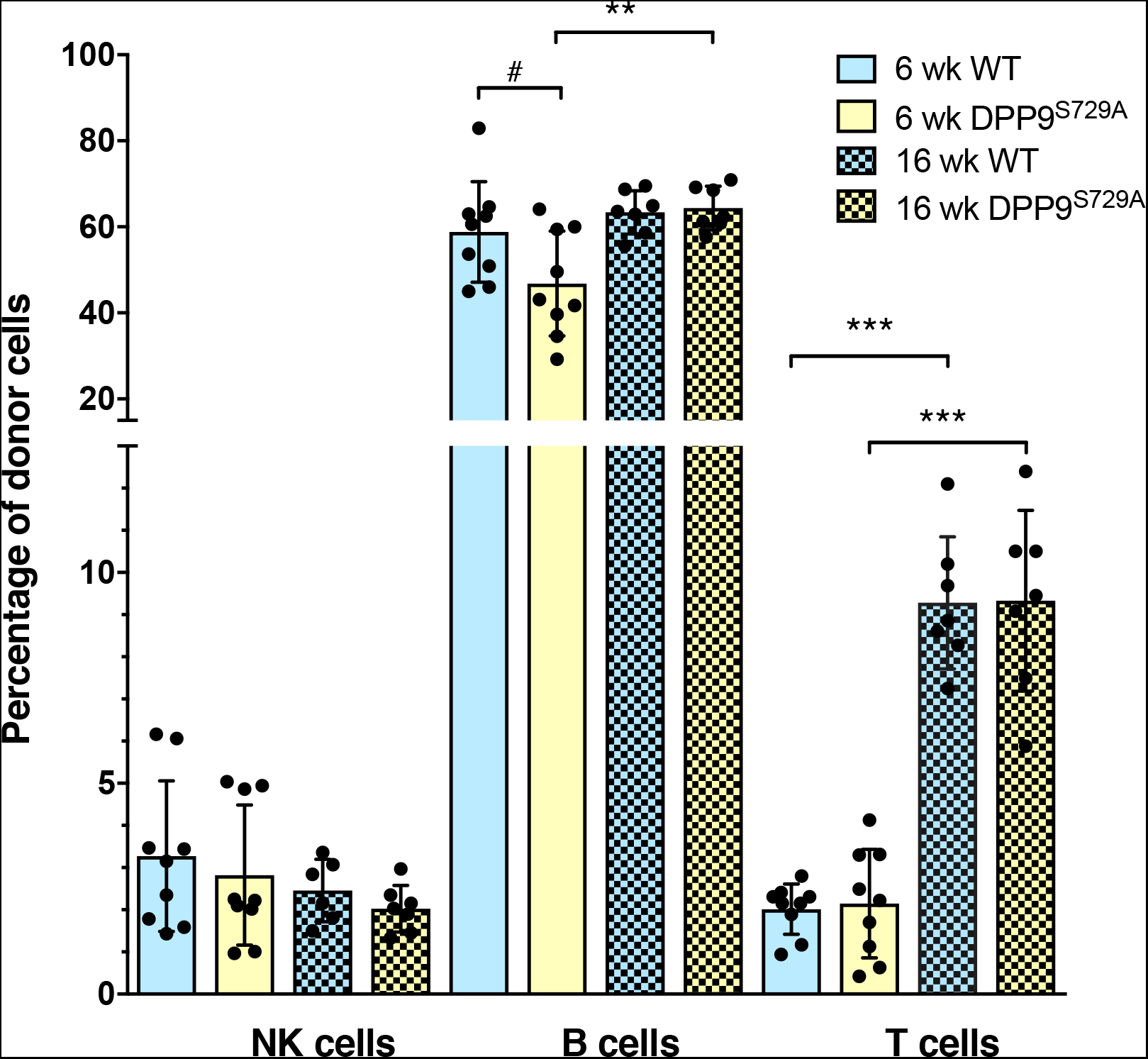
Myeloid and lymphoid cell phenotypes in peripheral blood of primary chimeric mice. Flow cytometry of peripheral blood from primary chimeric mice, showing total myeloid cells and neutrophils (A) and the major lymphoid cell phenotypes (B) as percentages of total donor leucocytes at 6 and 16 weeks post-transplantation. The proportions of myeloid cells and neutrophils were similar in DPP9^S729A^-origin mice and WT-origin mice. Myeloid cells and neutrophils were decreased at 16 weeks compared with 6 weeks for both groups (A). The proportion of B cells at 6 weeks was less in DPP9^S729A^-origin mice compared to WT-origin mice at 6 weeks, and compared to S729A-origin mice at 16 weeks (B). The proportion of T cells increased between 6 and 16 weeks in both DPP9^S729A^-origin mice and WT-origin mice (B). Mean, SD and individual data. n = 9 (6 weeks), n = 7 a 16 weeks. Significance; p < 0.05 (*), p < 0.01 (**), p < 0.001 (***).

There was a clear decrease in myeloid cells between 6 and 16 weeks post-transplantation (Figure 3A). This decrease was similar in both genotype-origins, and is consistent with changes that are known to occur between short-term and long-term regeneration after irradiation and donor engraftment ^37^. By 16 weeks, myeloid cell levels were comparable to those found in normal mouse peripheral blood ^36^. Neutrophils normally represent the largest percentage of myeloid cells, at approximately 30% in mouse. Neutrophils of the primary chimera were ~20% at 6 weeks and decreased to ~12% at 16 weeks post-transplantation, with no significant difference between genotype-origins (Figure 3A).

#### Identification of lymphoid cell types in primary chimeric mice

In these analyses, we assessed the major lymphoid lineages, namely CD3^+^ T cells, CD19^+^ B cells, and NK1.1^+^ natural killer (NK) cells. NK cells, which contribute to the early detection of virus-infected and cancer cells, are important for defense against pathogens and regain normal cell numbers and function within a month of transplantation ^28,38,39^. In the primary chimeric mice, no significant difference in NK cell numbers was observed between different genotype-origin groups or between 6 and 16 weeks post-transplantation (Figure 3B). These data suggest that once re-established in the peripheral blood, NK cell numbers were stable.

As reappearance of B cells is an early event occurring in the first few days after engraftment, the percentage of total B cells amongst the total lymphoid donor cells was expected to have reached a stable level by 6 weeks post-transplantation. This was the case for peripheral blood leukocytes at 6 weeks, although there were fewer B cells present in the DPP9^S729A^-origin cells compared to WT-origin cells (Figure 3B). At 16 weeks post-transplantation, the B cell numbers for both genotype-origin groups of mice were similar (Figure 3B) and consistent with normal peripheral blood B cell percentages ^36^.

Residual T cells of recipient origin, which is the cell type most resistant to irradiation, can be readily detectable within the first three weeks. Donor-derived T cells become readily detectable 4 to 5 weeks post-transplant ^29^. Since the primary chimeric mice received a lethal dose of myelo-ablative radiation, few residual recipient T cells were expected to survive. Concordantly, few donor T cells were detected 6 weeks post-transplantation (Figure 3B). T cell numbers significantly increased over time, reaching ~10% of peripheral blood leukocytes at 16 weeks post-transplantation. As T cells comprise about 20% of peripheral blood leukocytes in normal mice ^36^, full reconstitution of the T cell complement was perhaps not attained at 16 weeks post-transplantation in any of the mice. No significant difference in T cell numbers was observed between the WT and DPP9^S729A^ cell-origin recipient mice.

### Secondary chimeric mice of DPP9^S729A^-origin displayed an enhanced engraftment ability and an unimpaired capacity for self-renewal and proliferation

Successful long-term engraftment in a primary cell transplant recipient does not rule out defects in self-renewal capability. Hence, further serial transplants are often undertaken in studies of chimeras, in order to detect defects in HSC self-renewal capacity and the capacity for proliferation ^26^. Therefore, a second set of chimeric mice was generated, in which irradiated WT hosts received BM cells from primary chimeric mice along with BM cells from congenic PTPRC^A^ mice as competitor cells. As the result of the secondary engraftment was unknown before commencement of the experiment, competing donor cells and sub-lethal recipient irradiation were used to ensure successful cellular engraftment, immune regeneration and survival of the recipient mice ^27^.

Both mouse groups, which respectively received BM from the DPP9-WT-origin and DPP9^S729A^-origin primary chimeric mice, showed a similar pattern of weight loss and gain throughout 13 weeks of weight monitoring (Supplementary Figure 6). Return to start weight after initial weight loss was achieved in a much shorter time frame (about 2 weeks post-transplantation) than for the primary chimeric mice (4–5 weeks post-transplantation). This is consistent with the sub-lethal irradiation dose administered to the secondary mice whereas the primary mice received a higher-dose lethal myeloablative treatment.

#### Donor cell survival and replenishment in secondary chimeric mice

Irradiated mice were transplanted with DPP9^S729A^-origin and DPP9-WT-origin primary chimeric mouse BM cells along with PTPRC^A^ adult BM cells in the ratio of 60:40 (donor:competitor). As both competitor cells and recipient cells carried the CD45.1 allele, it was not possible to differentiate between these two sources of cells. Donor cells, however, all carried the CD45.2 allele and so could be clearly identified.

At both 6 and 16 weeks post-transplantation, the ratio of donor cells to competitor cells in WT-origin mice was 20:80 where the 80% includes residual recipient cells that survived irradiation (Figure 4A). This shows a dominance of competitor and residual recipient cells over donor cells, possibly due to increased numbers of residual recipient cells after sub-lethal irradiation. Interestingly, for the DPP9^S729A^-origin groups, at both time-points, the ratio of donor to competitor (with residual recipient) cells more closely aligned to the original transplanted cell ratio (40:60, Figure 4A), suggesting that the DPP9^S729A^-origin donor cells had an enhanced engraftment ability. The ratio of donor to competitor cells did not alter between 6 and 16 weeks post-transplantation. Proportions of individual cell populations are presented for each mouse (Figure 4B). Increased proportions of DPP9^S729A^-origin cells occurred across all lineages, suggesting better engraftment of HSC rather than improved competition by specific cell subsets.

**Figure 4:**
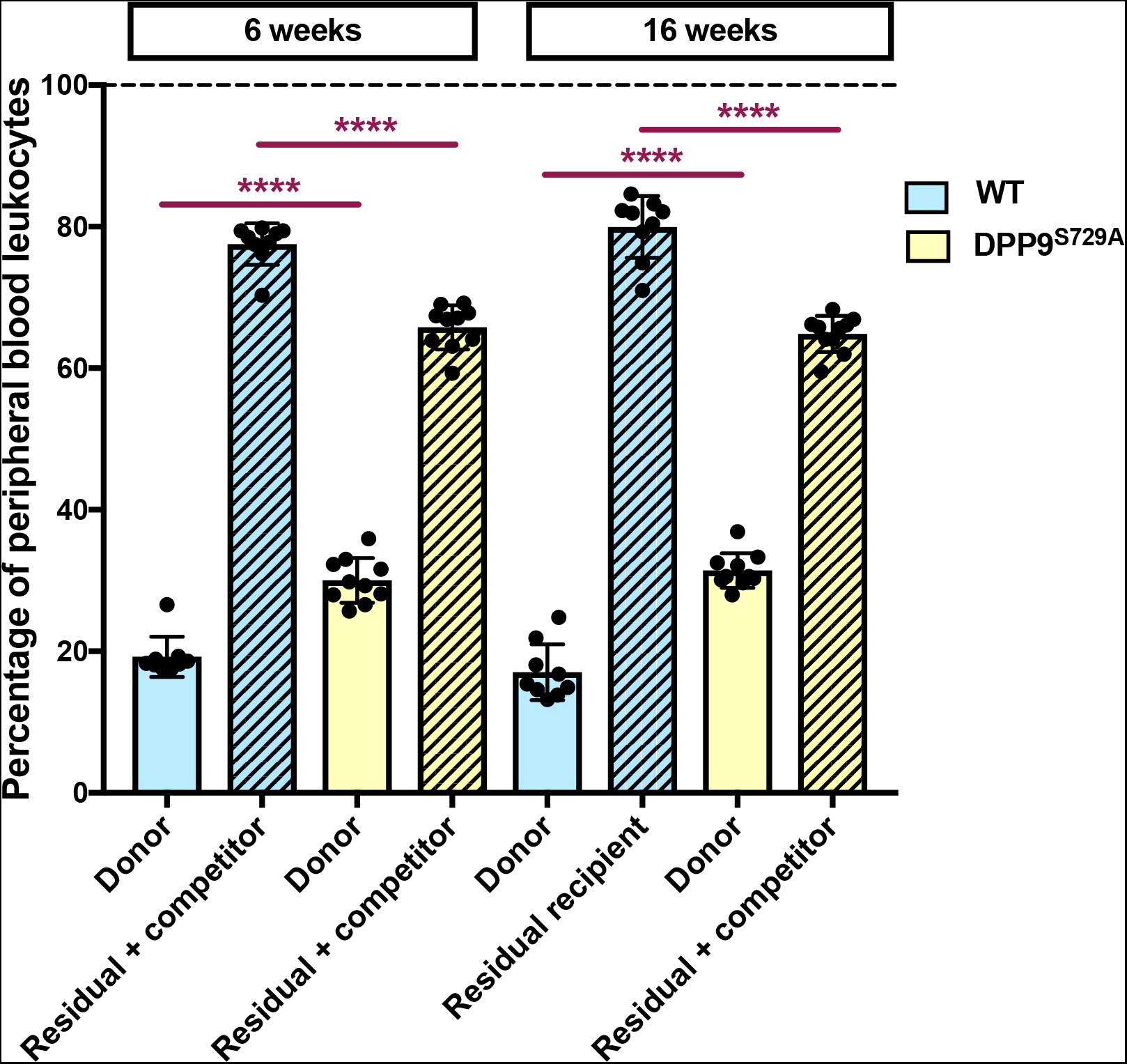
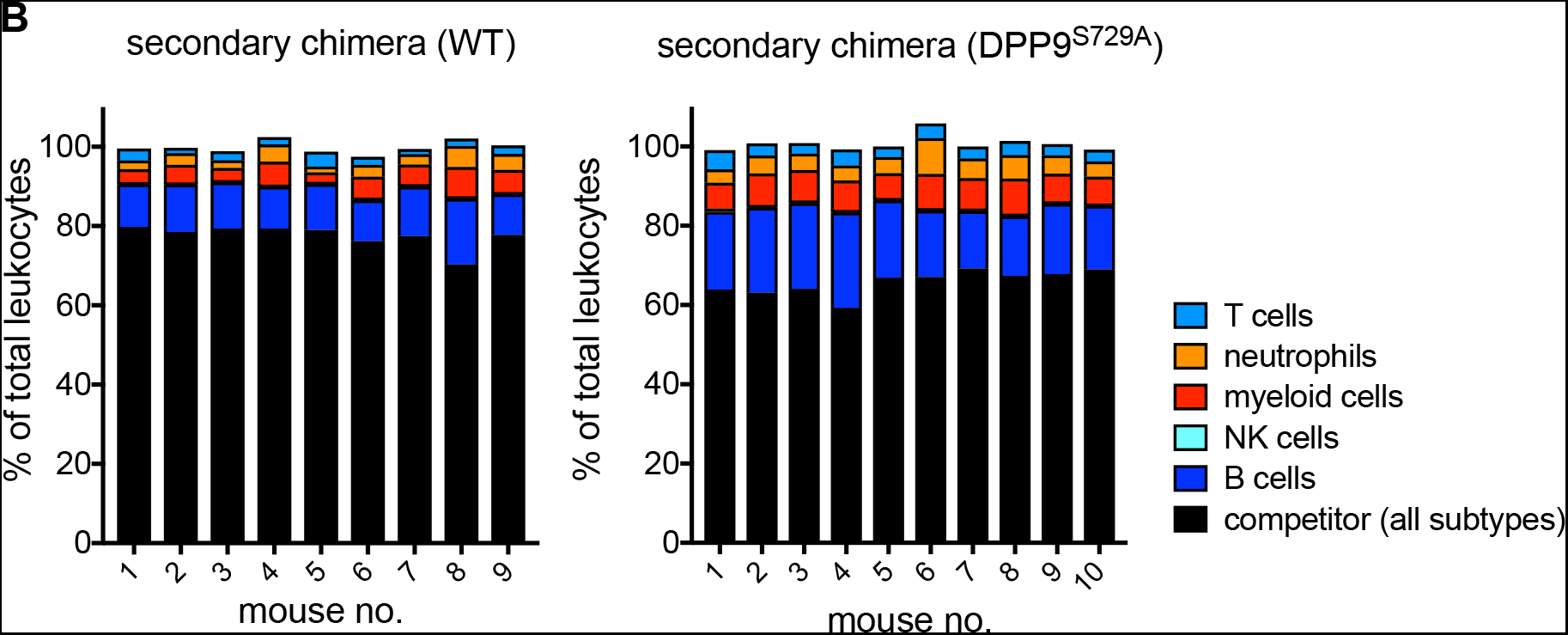
Proportions of donor and residual recipient cells of secondary chimeric mice. Irradiated mice were transplanted with DPP9^S729A^ or DPP9-WT adult bone marrow cells along with CD45.1^+^ *Ptprc^a^* adult BM cells at a ratio of 60:40 (donor:competitor). (A) The WT-origin group displayed a bias towards competitor cells over donor cells, whereas the DPP9^S729A^-origin group displayed no bias. All groups had a shortfall below 100% (dotted line), which represented outlier competitor cells and residual recipient cells that were gated out during analysis. Mean, SD and individual data. Significance; p < 0.0001 (****). (B) Proportions of leukocyte subpopulations for each mouse. WT-origin n = 9, DPP9^S729A^-origin n = 10, at 6 weeks and 16 weeks post-transplantation.

#### Identification of myeloid cell phenotypes in secondary chimeric mice

No differences in myeloid cell populations were observed between WT-origin and DPP9^S729A^-origin groups, at 6 or 16 weeks post-transplantation (Figure 5A). In a similar pattern to that seen in the primary chimeric mice, myeloid cell percentages showed a significant reduction from 6 to 16 weeks post-transplantation (Figure 5A). Myeloid cell percentages at 6 weeks were lower in the secondary chimera (~ 25%) compared to the primary chimera (~ 30%). Perhaps associated with this, the reduction in myeloid cell percentages at 16 weeks was less pronounced in the secondary than in the primary chimera. Similar to the primary chimera, approximately 60% of myeloid cells in each genotype were neutrophils at 6-weeks post-transplantation (Figure 5A). At 16 weeks, however, this proportion was approximately 50% for each genotype, and this difference was significant for DPP9^S729A^-origin cells. Perhaps at 16 weeks there was a less intense irradiation-induced inflammatory response in the secondary compared to the primary chimera, or a more rapid resolution of that response, such that at 16 weeks the myeloid cell compartment was normalised from a neutrophil-biased state to encompass other granulocyte cell types.

**Figure 5.**
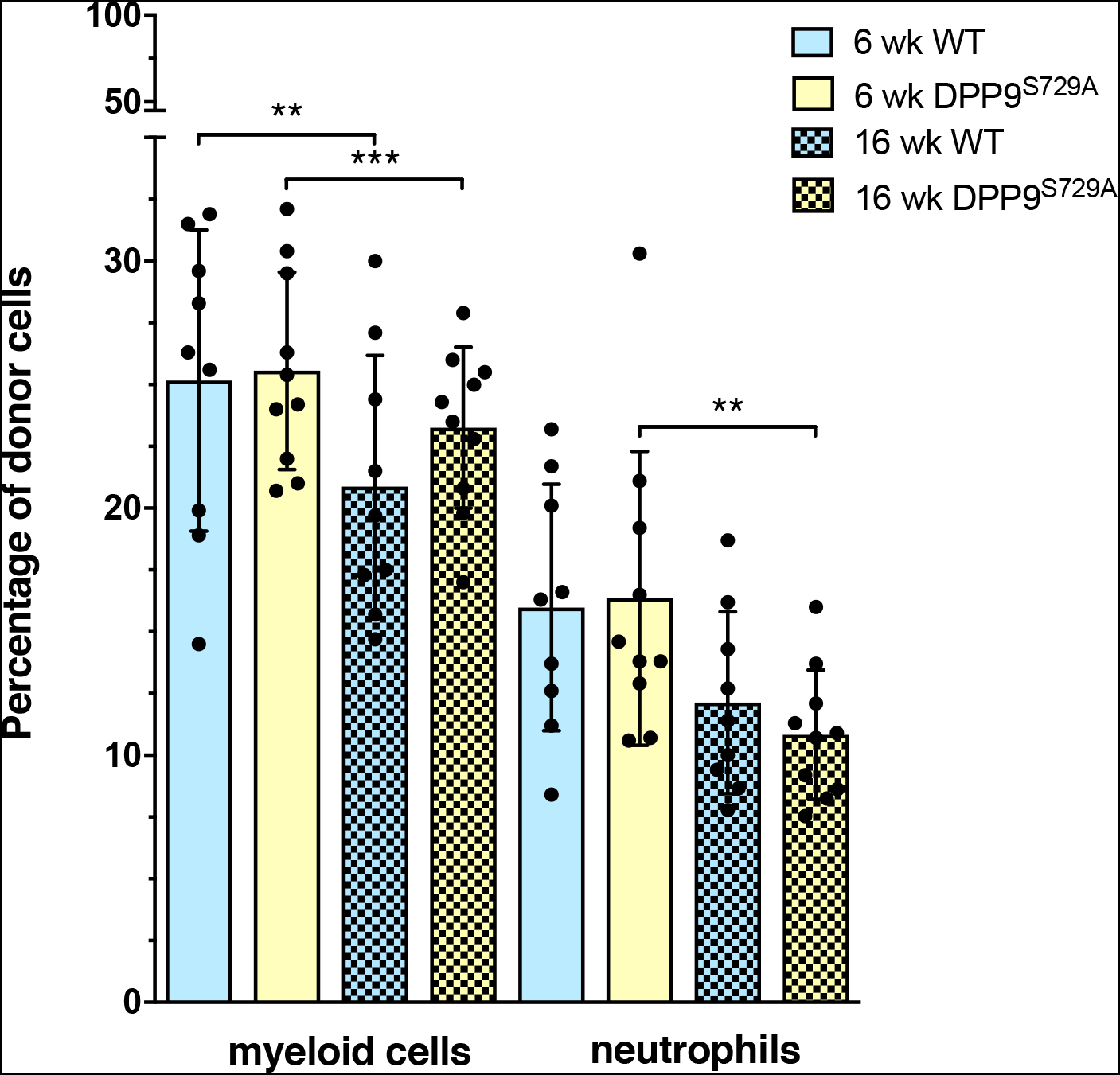
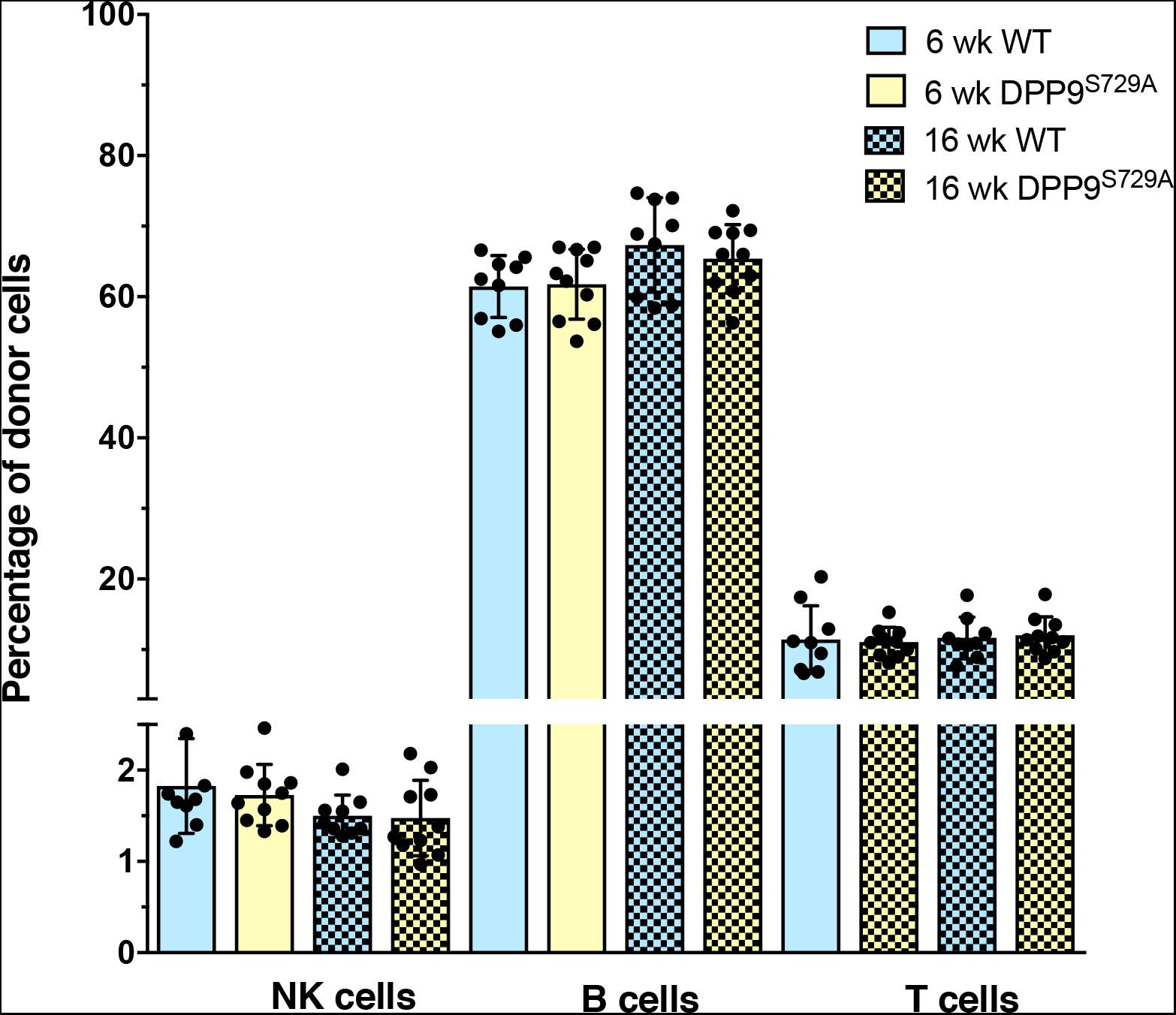
Myeloid and lymphoid cell phenotypes in peripheral blood of secondary chimeric mice. Flow cytometry of peripheral blood measured the percentages of total myeloid and neutrophil leukocytes (A) and of lymphoid lineage leukocytes (B) at 6 weeks and 16 weeks post-transplantation. Mean, SD and individual data, WT-origin n = 9, DPP9^S729A^-origin n = 10. Significance; p < 0.05 (*), p < 0.01 (**), p < 0.001 (***).

#### Identification of lymphoid cell phenotypes in secondary chimeric mice

In secondary chimeric mice, the main lymphoid lineage cell subsets were present and were not significantly different in abundance between genotypes (Figure 5B). As with the primary chimera, the B cell percentages were consistent with those normally observed in mouse peripheral blood leukocytes ^36^. In contrast to the primary chimeric mice, T cell percentages were approximately 11% at both 6 and 16 weeks post-transplantation in each chimera. This was less than the expected level of 20% T cells in peripheral blood leukocytes.

Considering the overall peripheral blood leukocytes of both myeloid and lymphoid lineage, the trend in the primary chimeric mice from 6 to 16-weeks post-transplant was towards a greater proportion of lymphoid cells (Figure 6A). This is consistent with a change from unlimited self-renewal ability by HSC towards a limited self-renewal ability as the regenerating immune system changes from short term to long term reconstitution ^40^. For the secondary chimeras, the peripheral blood leukocytes of both myeloid and lymphoid lineages were similar at 6 and 16 weeks post-transplant, with a strong bias towards the lymphoid lineages (Figure 6B).

**Figure 6:**
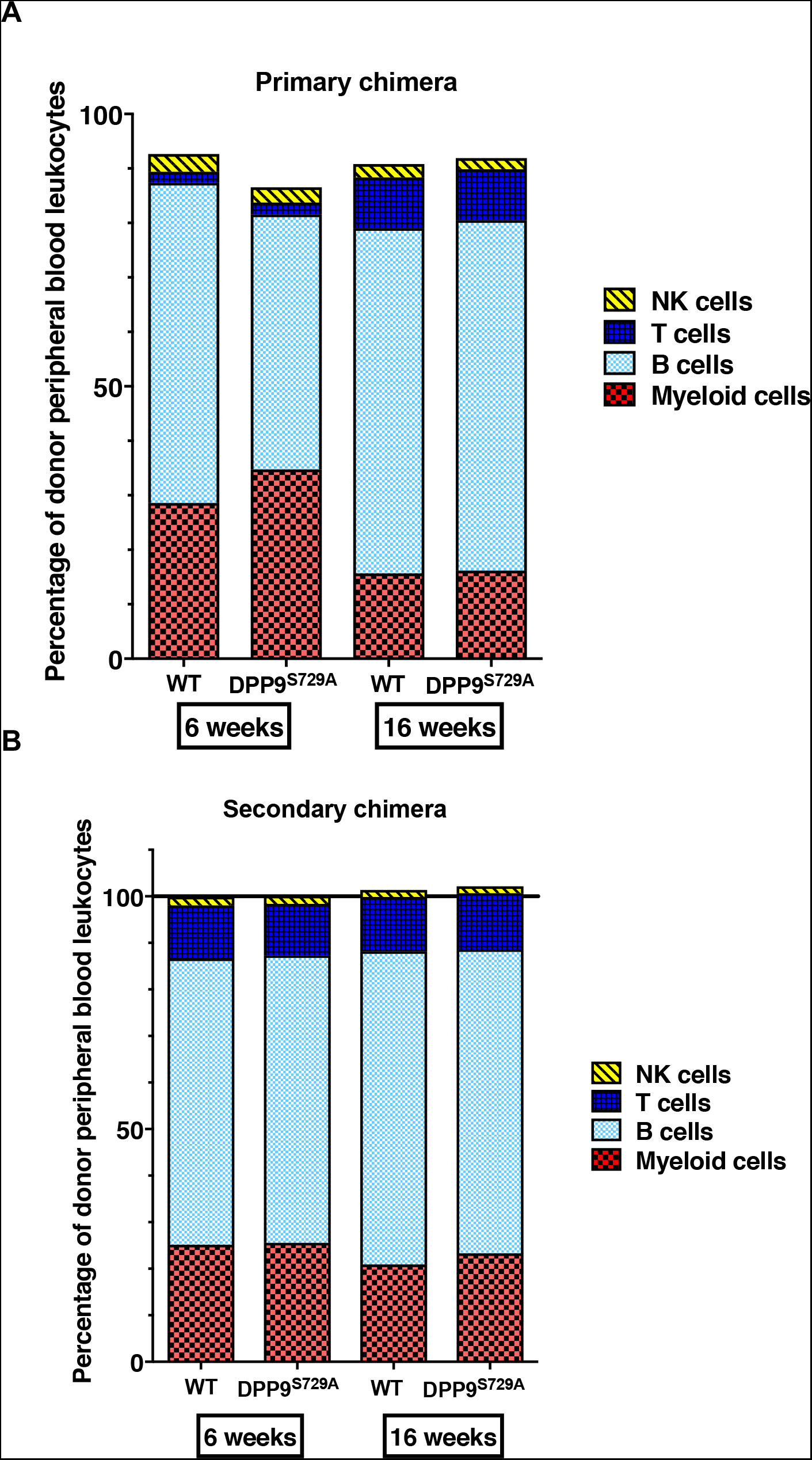
Enumeration of donor peripheral blood leukocytes of primary and secondary chimeric mice. Phenotypic analyses of total peripheral blood leukocytes (PBL) of donor origin. Between 6 and 16 weeks after fetal liver cell transplant, both WT-origin and DPP9^S729A^-origin mice increased the proportion of lymphoid lineage cells in PBL (A). The proportions of lymphoid and myeloid lineage cells were comparable between 6 and 16 weeks after bone marrow cell transplant of WT-origin or DPP9^S729A^-origin cells (B). The total percentage of cells were more than 100% (designated by the black line) at 16 weeks due to the univariate analysis used by the FlowJo software to preserve relative relationships between groups.

[BEN: See my comment above. It is also putting discussion material into the results section.]

## Discussion

We created chimeric mice using fetal and adult donor cells with both normal and absent DPP9 enzymatic activity. We showed that a lack of DPP9 enzymatic activity in HSC does not impair their ability to engraft in irradiated mice or to repopulate all major immune cell types in the regenerated immune system in primary or secondary chimeras. Interestingly, secondary chimeric mice were efficiently engrafted by blood immune cell precursors lacking DPP9 enzymatic activity.

Fetal liver cells of DPP9-WT and DPP9^S729A^ origin consisted of the same proportions of erythroid and non-erythroid lineage cells. This provided a common starting point for creation of the chimeric mice and for assessment and quantification of HSC numbers and functionality.

Our data broadly concord with data from an independently-derived DPP9^S729A^ mouse ^41^. That study examined only primary chimeras and only at 6 weeks and used ED 17.5 rather than ED 13.5 fetal liver. Importantly, the data from both that study and our study are consistent with outcomes current knowledge of normal engraftment by fetal liver cells ^36,37,39^, and so equally suggest that lacking DPP9 does not impair engraftment. The reconstitution of the primary chimeric mice was very high at both 6 and 16 weeks post-transplant, with only small percentages of residual recipient cells present. Taken together, there was no detectable impairment in the ability of DPP9^S729A^ HSC to engraft in the primary chimera. For the secondary chimeric mice, with an original transplant ratio of 40:60 of donor to competitor + residual cells, the WT-origin group had a higher ratio (20: 80) whilst the DPP9^S729A^-origin group had a similar ratio to that of the original transplant. Thus, the DPP9^S729A^-origin donor cells may possess an enhanced engraftment ability in the secondary chimera. Ablation of the DPP9 – related enzyme DPP4 enhances HSC engraftment in irradiated mice ^21^, so this might be a general property of inhibitors of this enzyme family.

Having confirmed effective engraftment of donor cells, we then determined the contribution of hematopoietic cell lineages to the development and repopulation of the reconstituting immune system. At 6 and 16 weeks post-transplantation, in both the primary and secondary chimeric mice, all myeloid and lymphoid cell types considered in this study were present in both genotype-origin groups. Therefore, the progenitor cell pool was intact and both the primary and secondary HSC showed no defect in self-renewal or proliferation capability with the absence of DPP9 enzymatic activity.

The total myeloid lineage cell numbers did not differ with genotype. NK cell numbers were stable regardless of genotype. T cell reconstitution showed no difference between genotype.

B cells were present in comparable levels in both the primary and secondary chimeras, however, in the primary chimeric mice, fewer B cells in the DPP9^S729A^-origin group were seen compared to the WT-origin group at 6 weeks but not at 16 weeks. This may reflect a delay in B cell re-establishment in the DPP9^S729A^ group. Taken together, these results indicate that the lack of DPP9 enzymatic activity in hematopoietic cell chimeras affected granulocyte subsets and B cells but not NK cells or T cells. DPP9 inhibition stabilizes Syk, thereby modulating Syk signaling in B cells ^8^. Thus, DPP9 enzyme activity might have a role in B cell function that influences differentiation and/or proliferation in the absence of antigen stimulation.

In summary, this work provides insights into the roles of DPP9 enzymatic activity in immune and hematopoietic systems and contributes to the understanding the DPP9 protease in biological systems.

## Acknowledgements

This work was supported by the National Health and Medical Research Council of Australia Project Grant 512282 (MDG) and Program Grant 571408 (GWM), The Rebecca L. Cooper Medical Research Foundation, a University of Sydney Equipment Grant, Sydney Medical School Foundation/Francis M. Hooper Scholarship for Medical Research through the University of Sydney (MGG), and an Australian Postgraduate Award (HEZ). The authors thank Jeff Crosbie for mice irradiation. The authors acknowledge the animal facility and the flow cytometry facility at the Centenary Institute for the scientific and technical assistance.

## Supplementary Figures S1 to S5

**Figure S1:**
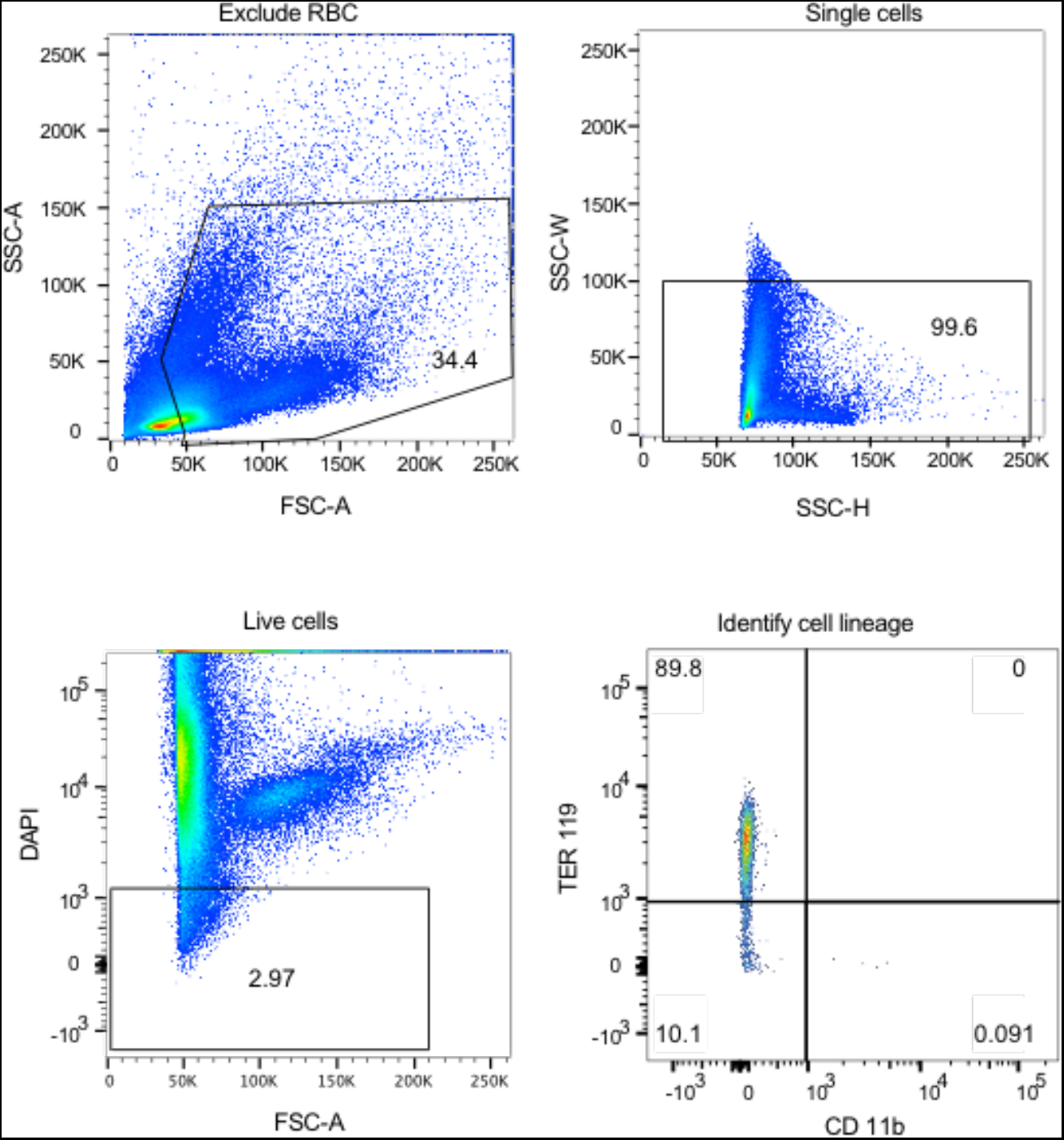
Flow cytometry gating strategy for fetal liver cells. Gating excluded red blood cells remaining after lysis and then excluded cell doublets and dead cells (DAPI). Live cells were then gated to identify TER-119 positive and CD11b positive cells. TER-119 positive cells indicate cells of the erythroid lineage while CD11b positive cells indicate cells of the myeloid lineage. This representative flow cytometry data derives from WT cells.

**Figure S2:**
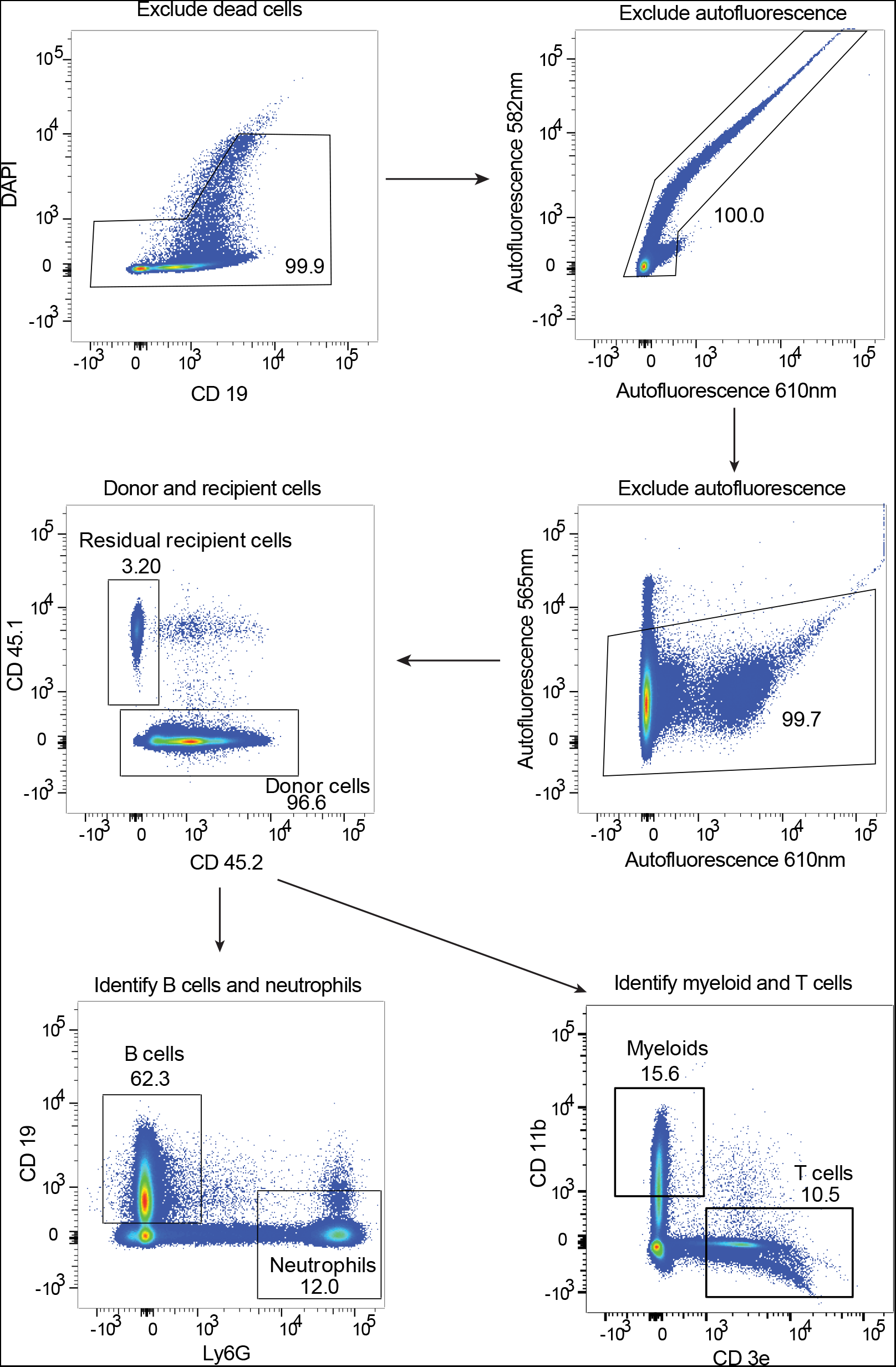
Flow cytometry gating strategy for donor and recipient cell identification. Live/dead cells in PBL were gated using DAPI staining and then autofluorescent cells were excluded at 565nm, 582nm and 610nm. Then CD45.1 positive recipient cells and CD45.2 positive donor cells were identified. CD11b and CD3e were markers of myeloid cells and T cells, respectively. Ly6G and CD19 were markers of neutrophils and B cells, respectively. Representative data from DPP9-WT-origin chimeric mice 16 weeks after irradiation.

**Figure S3:**
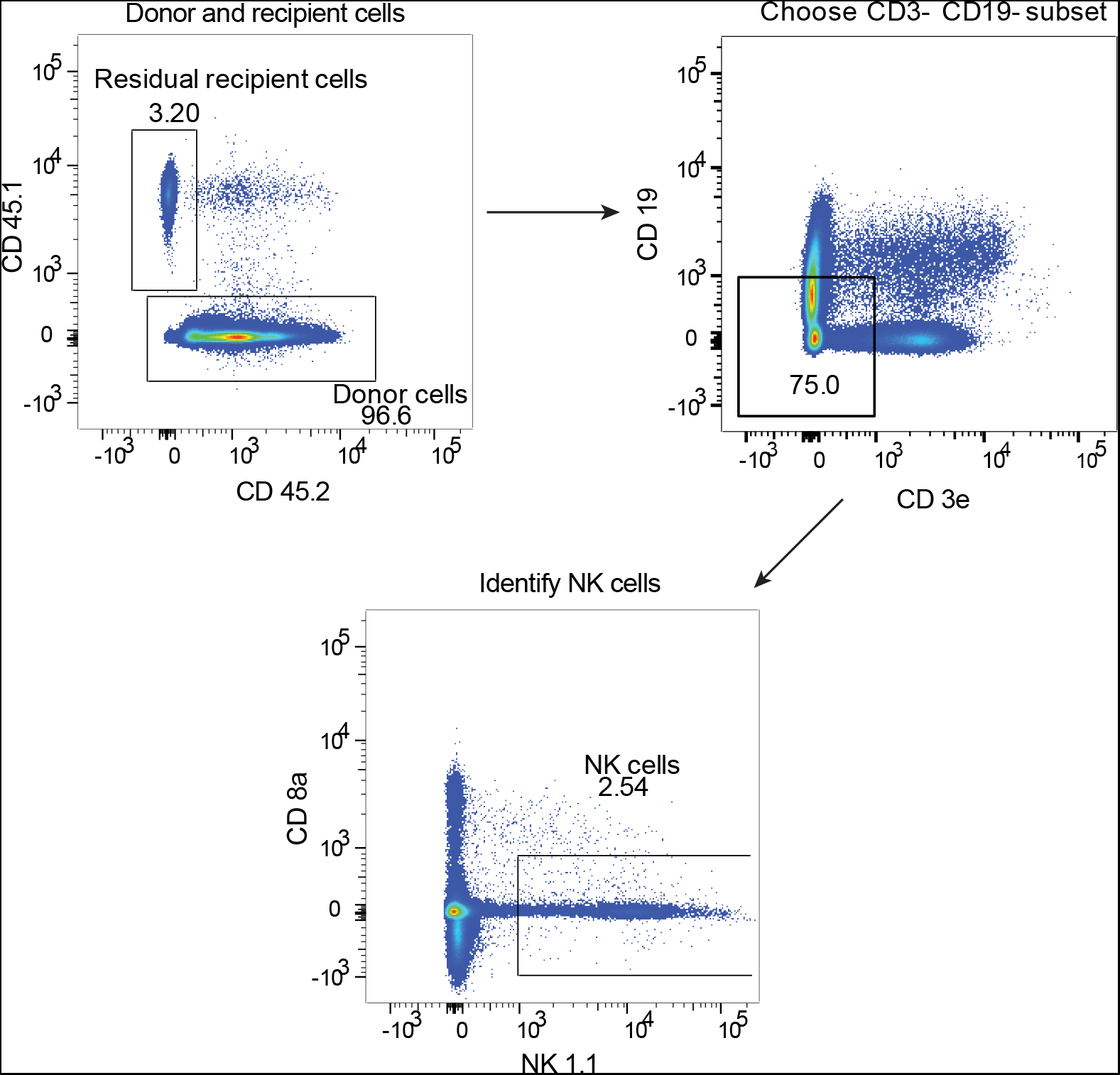
Flow cytometry gating strategy for identifying NK cells. CD45.2 positive donor cells were identified in PBL as described in Figure S2. The markers CD3e and CD19 were used to identify the CD3e^−^CD19^−^ cell subset that includes NK cells. The markers CD8a and NK1.1 were then used to identify NK cells. Representative data from a DPP9-WT-origin chimeric mouse 16 weeks after irradiation and transplantation.

**Figure S4:**
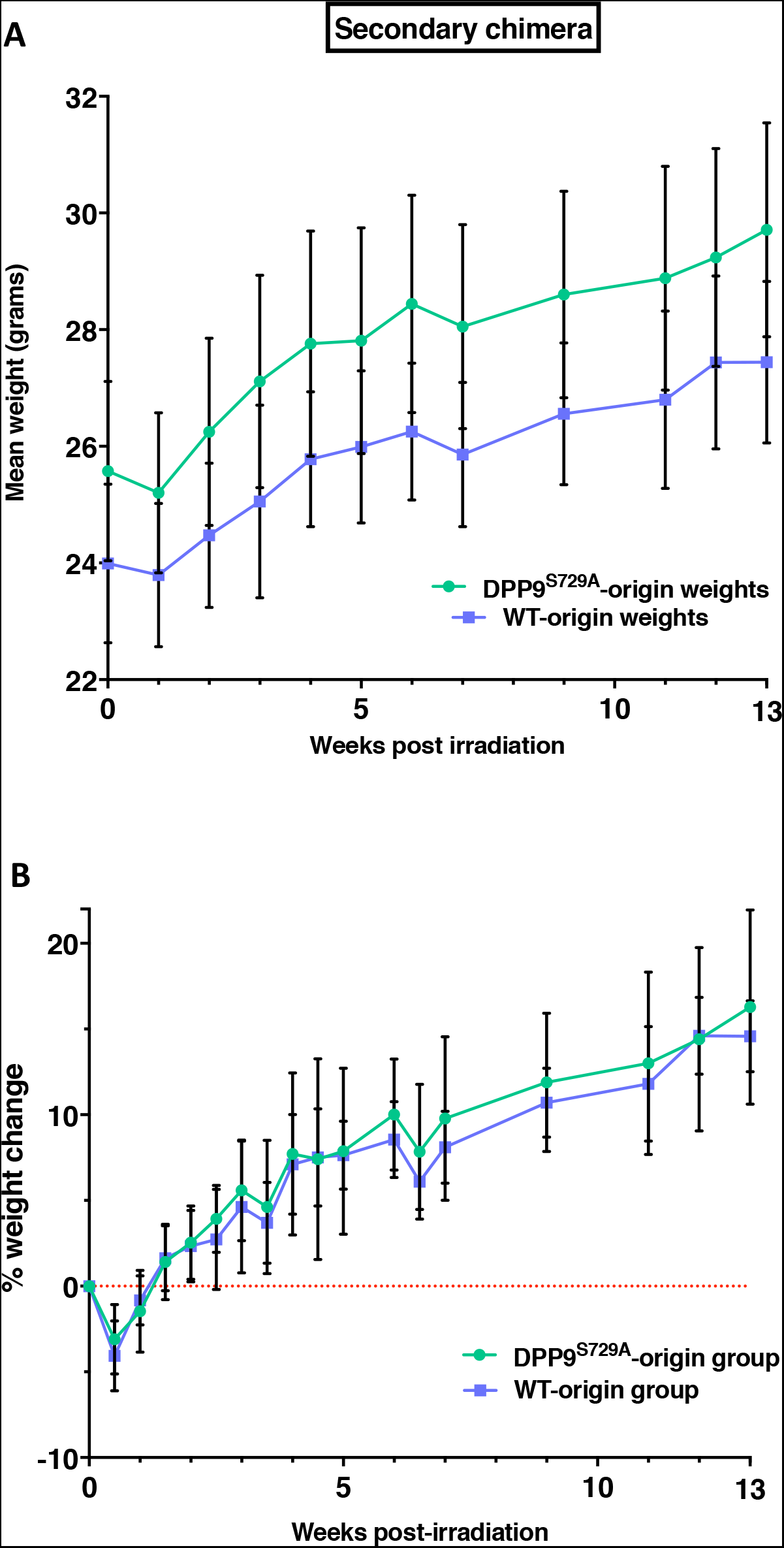
Body weight: primary chimeras. Irradiated mice inoculated with DPP9^S729A^ or DPP9-WT fetal liver cells were monitored for weight loss as a gross indicator of dysfunctional or failed immune regeneration. All irradiated mice exhibited an initial weight loss for several days before gradual weight increase back to the original start weight at around 4–5 weeks post-transplantation. Mean body weight of mice (**A**), and mean percentage weight change where 0 (red dotted line) represents the body weight immediately before irradiation (**B**). n = 7 mice per group.

**Figure S5:**
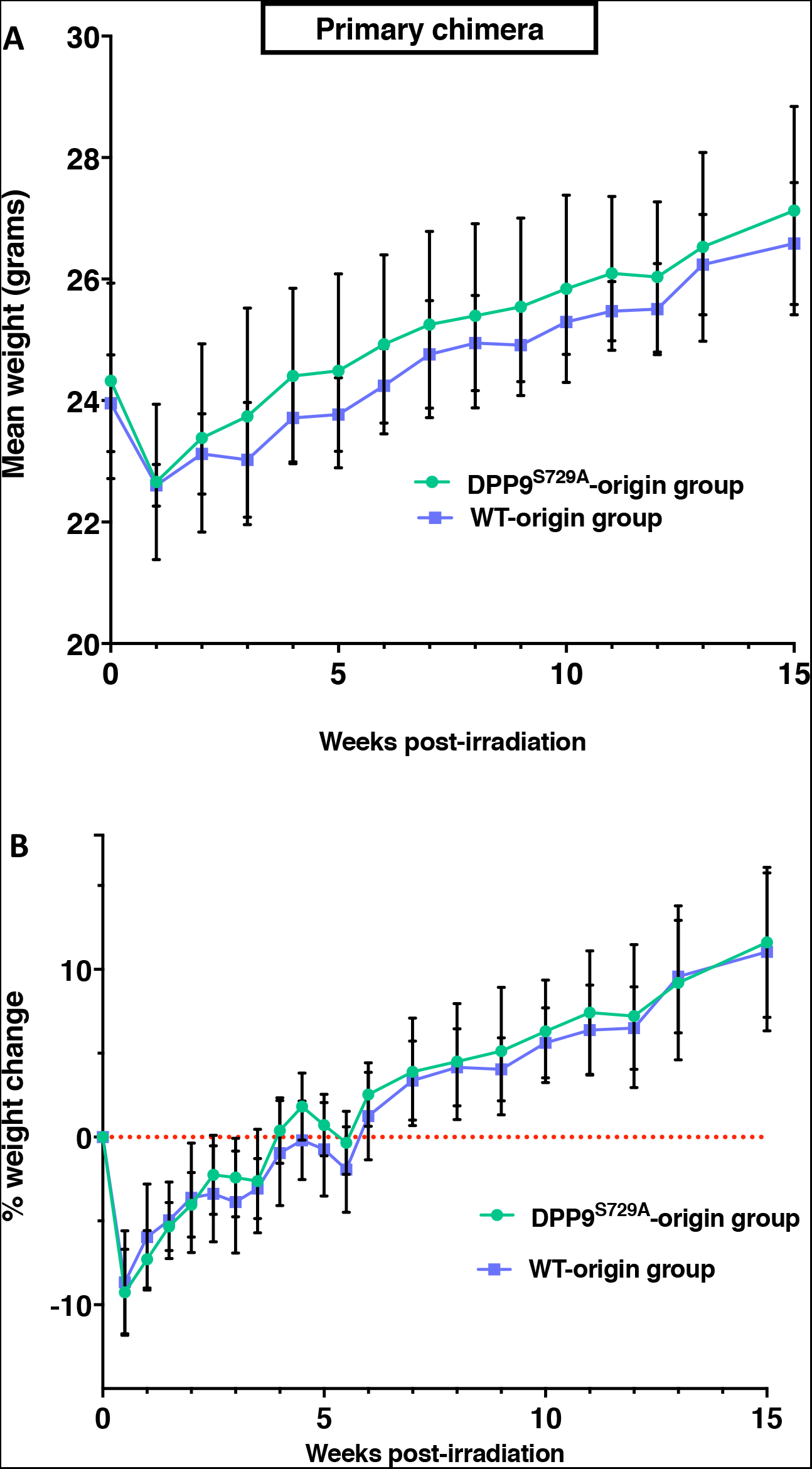
Body weight: secondary chimeras. Irradiated mice inoculated with DPP9^S729A^ (n = 10) or DPP9-WT (n = 9) bone marrow cells along with PTPRC^A^ WT bone marrow cells were monitored for weight loss as a gross indicator of dysfunctional or failed immune regeneration. All irradiated mice displayed an initial weight loss for several days before gradual weight increase, returning to the original weight by 2 weeks after irradiation. Mean body weight of mice (**A**), and mean percentage weight change where 0 (red dotted line) represents the body weight immediately before irradiation (**B**).

## Abbreviations

BM: : bone marrow
DAPI: : 4’,6-diamidino-2-phenylindole
DMSO: : dimethyl sulfoxide
DPP: : dipeptidyl peptidase
ED: : embryonic day
EDTA: : ethylenediaminetetraacetic acid
FACS: : fluorescence-activated cell sorting
FCS: : fetal calf serum
HBSS: , Hank’s balance salt solution
HSC: : hematopoietic stem cells
NK: : natural killer
WT: : wild-type

